# The Diversity and Ubiquity of Antibiotic Resistance Genes in Finfish Culture Ponds in Bangladesh

**DOI:** 10.1101/2022.09.14.507951

**Authors:** Ashley G. Bell, Kelly Thornber, Dominique L. Chaput, Neaz A. Hasan, Md. Mehedi Alam, Mohammad Mahfujul Haque, Jo Cable, Ben Temperton, Charles R. Tyler

## Abstract

In Bangladesh, fish provide over 60% of animal-source food with 56.2% of this coming from aquaculture produced predominantly in rural freshwater ponds. Increasing demand for fish products is driving intensification and resulting in higher disease prevalence, posing a risk to food security. Biosecurity is often absent in rural aquaculture practices in Bangladesh and antibiotics are commonly used to treat and prevent disease outbreaks. Antibiotics are often administered incorrectly - a key factor associated with the development of antimicrobial resistance (AMR). AMR can be disseminated rapidly within microbial ecosystems via mobile genetic elements, posing a risk for humans and animals infected with AMR pathogens as treatments with antibiotics become ineffective. Early AMR detection and understanding of the spread of antimicrobial resistant genes (ARGs) in rural aquaculture practices is critical for both food security and human health protection. Here, we apply a metagenomic approach to assess the ARG composition in pond water from six finfish (tilapia and pangasius) farms in the Mymensingh division of North-central Bangladesh. We found microbial communities within the ponds had similar alpha and beta diversities, with multiple ARGs predicted to confer resistance to eighteen different classes of antimicrobials. The most common ARGs conferred resistance to aminoglycosides and sulphonamides and were present in taxa associated with both fish and human pathogens. This ARG diversity potentially confers resistance to a wide variety of antibiotic classes and questions the effectiveness of current and future treatment of diseases with antibiotics in earthen aquaculture ponds. The microbial and ARG compositions between fish ponds within each farm were similar, which may relate to parallels in farming practices creating similar microbial selection pressures and thus comparable microbial populations. Without a more controlled approach towards antibiotic usage, will inevitably further exacerbate the challenges in treating and preventing disease outbreaks as aquaculture production intensifies in Bangladesh.

**Highlights:**

- ARGs in Bangladesh rural fishponds indicate resistance to 18 different antibiotics
- The most common AMR were to aminoglycosides and sulphonamides
- ARGs were present in plasmids and taxa-associated pathogens
- Farming practices strongly influence microbial and ARG compositions
- Identified ARGs question antibiotic treatment of disease in rural aquaculture

## 1. Introduction

Antibiotics from multiple anthropogenic activities enter freshwater systems via various sources including wastewater effluent, surface runoff from agriculture, and aquaculture (Cabello, 2006; Czekalski et al., 2014; Dolliver & Gupta, 2008; Michael et al., 2013; Schar et al., 2020). This has contributed to the growing problem of antimicrobial resistance (AMR) through the reduction of antibiotic efficacy, as pathogens acquire antimicrobial resistance genes (ARGs). Much of the AMR problem has been attributed to the increasing burden of infectious disease, coupled with our misuse of antibiotics in the absence of effective infectious disease control, and limited alternative treatment therapies (Ventola, 2015). Unless addressed, it has been estimated that the AMR crisis will cost the world economy USD 100 trillion by 2050 and lead to the loss of 10 million lives annually (O’ Neil, 2014). The implementation of national action plans outlining strategies to reduce or mitigate AMR risks is key to combating rising AMR rates (European Commission, 2021; WHO, 2017). A key part of these action plans is the reduction of AMR risks associated with food production, where around two-thirds of all antibiotics are used (Chang et al., 2015; Hollis & Ahmed, 2013; Kennedy, 2013).

Aquaculture has been identified as a key contributor to sustainable protein production for a growing global human population (UN General Assembly, 2015). The majority (89% in 2019) of aquaculture occurs in Low- and Middle-Income Countries (LMICs) within Asia. This includes Bangladesh, whose aquaculture industry contributed 3% of global fish production by weight (FAO, 2020), with an annual growth rate of roughly 10% over the last decade (FAO, 2020; Hossain et al., 2015; Shamsuzzaman et al., 2020). Over half (56.2%) of Bangladesh’s fish production now comes from aquaculture (FAO, 2020), and the industry is undergoing rapid intensification to meet domestic economic growth targets and rising global demand for animal products (Schar et al., 2018). Traditional and rural aquaculture farms comprise small-scale earthen embankment ponds that are closely connected to their local environment. However, with poor access to the infrastructure or resources to support bio-secure intensification, the industry is experiencing increasing levels of disease (Henriksson et al., 2018), putting the livelihoods of the 15 million people at risk who depend upon Bangladesh fisheries (DoF, 2019).

Antibiotics are commonly administered to ponds to treat and prevent disease, and reports suggest they continue to be used as growth promoters (Chowdhury et al., 2021; Rousham et al., 2019). The socioeconomic drivers of antibiotic overuse in Bangladesh aquaculture are complex and multifactorial, with farmers facing challenges from increasing disease burden, low and unpredictable market prices, poor source water quality and little access to training on sustainable practices and disease management (Abu Kawsar et al., 2019). Antibiotics enter pond environments from their (illegal) use in commercial fish feeds and/or prophylactically, where they are added to prevent spoiling, or through contamination of pond water with antibiotic-treated human and animal waste from the wider pond environment. Antibiotics are cheap and readily available over the counter without a prescription in Bangladesh (Rousham et al., 2019; Schar et al., 2020), and reports suggest that they are commonly applied to ponds at doses higher than those recommended and without completion of the full course of treatment to treat diseases (Abu Kawsar et al., 2019; Ali et al., 2016; Jahan et al., 2015), factors that are associated with the development of AMR (Fair & Tor, 2014). Water used on fish farms is generally discharged without treatment into the same supplying water body leading to higher antibiotic concentrations at downstream sites and the intakes for other fish farms (Belton et al., 2011; Jahan et al., 2015; Thompson et al., 2000). Antibiotics are the most common (80.85%) veterinary drugs used in fish farming in Bangladesh (Faruk et al., 2021) and their use should follow national and international guidelines as well as codex regulations (Ababouch, 2014). However, there is no legislation in place regulating the use of veterinary drugs in aquaculture (FAO, 2022). These factors, combined with poor diagnostics, limited veterinary services and lack of enforced regulation on antibiotic usage have been highlighted as major causes of reduced antibiotic efficacy in fish ponds (Iskandar et al., 2020).

Recent studies now evidence the presence of ARGs in Bangladesh aquaculture environments (Hemamalini et al., 2022; A. Hossain et al., 2022; Lassen et al., 2021; Thornber et al., 2020). Within Bangladeshi fish ponds, untreated wastewater often containing AMR microbes are also used for other crop production, bathing, washing clothes and food preparation, which facilitates the passage of ARGs via bacteria into humans (Amin et al., 2019). Chronic antibiotic exposure in humans, livestock and wild animals, including freshwater fish, has been shown to result in an increased incidence of difficult-to-treat infectious diseases due to the development of AMR (Ikhimiukor et al., 2022; Lassen et al., 2021; Manyi-Loh et al., 2018; Marshall & Levy, 2011). Although the presence of ARGs within non-pathogenic organisms may not be a direct risk to human or animal health, chronic exposure of microbes to antibiotics creates a positive selection pressure for the acquisition of ARGs. ARG acquisition occurs through various routes, including horizontal gene transfer, which in turn increases the number of ARGs within a local system (Davies & Davies, 2010; Kraemer et al., 2019; Lerminiaux & Cameron, 2018). This acquisition of ARGs on mobile genetic elements, such as plasmids, which are readily transferred between bacterial species, contributes to rapid dissemination within microbial ecosystems and into pathogens (Partridge et al., 2018; Von Wintersdorff et al., 2016).

The high potential for the emergence and transmission of AMR from LMIC aquaculture environments has been recently recognised (Ikhimiukor et al., 2022; Koutsoumanis et al., 2021; Thornber et al., 2020), with calls for greater monitoring and surveillance of resistant pathogens and mobile genetic elements in these contexts (European Commission, 2021; FAO, 2016). Traditional AMR surveillance and monitoring is labour intensive, requiring culturing or quantifying ARGs through qPCR reactions, and is becoming increasingly expensive with large and growing ARG databases. In Bangladesh, there has been a recent increase in the use of shotgun metagenomic methods (Majeed et al., 2021; McInnes et al., 2021), which when sequenced allow for unlimited re-analyse against newly discovered ARGs and is not biased against the minority of microorganisms that are culturable (Steen et al., 2019). Taxonomic identification of sequences allows the linkage of ARGs with taxa of interest such as pathogens and mobile genetic elements whilst maintaining important metrics such as ARG abundance within a metagenome. Metagenomic studies in Bangladesh have linked the abundance of ARGs in rivers and fish ponds with bacteria from the human gut suggesting untreated wastewater as a source of AMR (McInnes et al., 2021) and applied metagenomic data for AMR surveillance (Majeed et al., 2021). ARG prevalence and composition have also been shown to be associated with geography and feed type in fish ponds in Bangladesh (Lassen et al., 2021).

In this study, we sought to examine how microbial assemblages, ARG abundance and diversity differed between seasons, crop type (fish) and geographical region in Bangladesh aquaculture pond water samples using a metagenomic shotgun DNA sequencing approach. Between 2017 and 2018, pond water was collected from six different fish farms at four seasonal time points containing either Nile tilapia (Oreochromis niloticus) or pangasius (Pangasianodon hypophthalmus) in the Mymensingh division (region) of Northern Bangladesh. Mymensingh is the highest pond-based fish-producing district in Bangladesh with an associated high usage of pharmaceuticals (including antibiotics). We identified the diversity and taxonomy of microorganisms that encode ARGs within Bangladeshi fish ponds to better understand their proliferation as plasmids and their presence within putative pathogens. We found no differences in the microbial community compositions or ARG profiles across all sample points suggesting that similarities in farming practices, including antibiotic usage, may be more influential than environmental factors in shaping the microbial community compositions and ARG profiles in these pond systems.

## 2. Methods

### 2.1. Sample locations and pond cultures

We applied metagenomic sequencing to investigate the microbial and ARG composition and diversity in finfish ponds at six different fish farms in the Mymensingh division in North-central Bangladesh. In a previous study, we used a smaller metagenome data analysis from these pond samples to support the building of a framework to better understand the risks of antimicrobial resistance in aquatic food-producing environments in Bangladesh (Thornber et al., in review). Here were expanded our analysis significantly to assess ARG types, abundance and multi-resistance genes in these ponds to assess for differences across space and time (over seasons). The six different fish farms sampled for this study were spread over five different villages and three Upazilas (subdistricts), with the farm’s main finfish crop being either Nile tilapia (Oreochromis niloticus) or pangasius (Pangasianodon hypophthalmus) (Figure 1B) which are among top three commercially important fish species in Bangladesh (Rahman et al., 2021). Fish ponds were rural earthen ponds enclosed by dykes that varied in size from 1200 m^2^ and 16000 m^2^ (mean 4000 m^2^, median 2500 m^2^) and between 1 m and 1.5 m in depth. Farmers at each farm were retrospectively interviewed in 2021, as described in (Thornber et al., in review) to ascertain the pond conditions in which the fish were reared. Ponds were topped up, on average weekly, with a mixture of rainwater and groundwater sources depending on rainfall conditions and seasons. Tilapia are harvested and the ponds are restocked every three to four months, whereas pangasius has an annual cropping cycle. On a rotational basis, ponds are dried, limed to reduce acidity, disinfected using bleach and prepared for re-stocking crop (fish) for the next season. In the off-seasons, every two to three years, sludge from the pond bottom is removed. Fish are generally fed a mixture of sinking or floating commercial feed, although some farmers also supplement the fish diet using their own farm food sources. A variety of antibiotics are administered before or during re-stocking events by mixing the antibiotic into the fish feed and/or water. A wide variety of antibiotic products are used, with the most common antibiotic classes being diaminopyrimidines, sulphonamides and fluoroquinolones.

**Figure 1.**
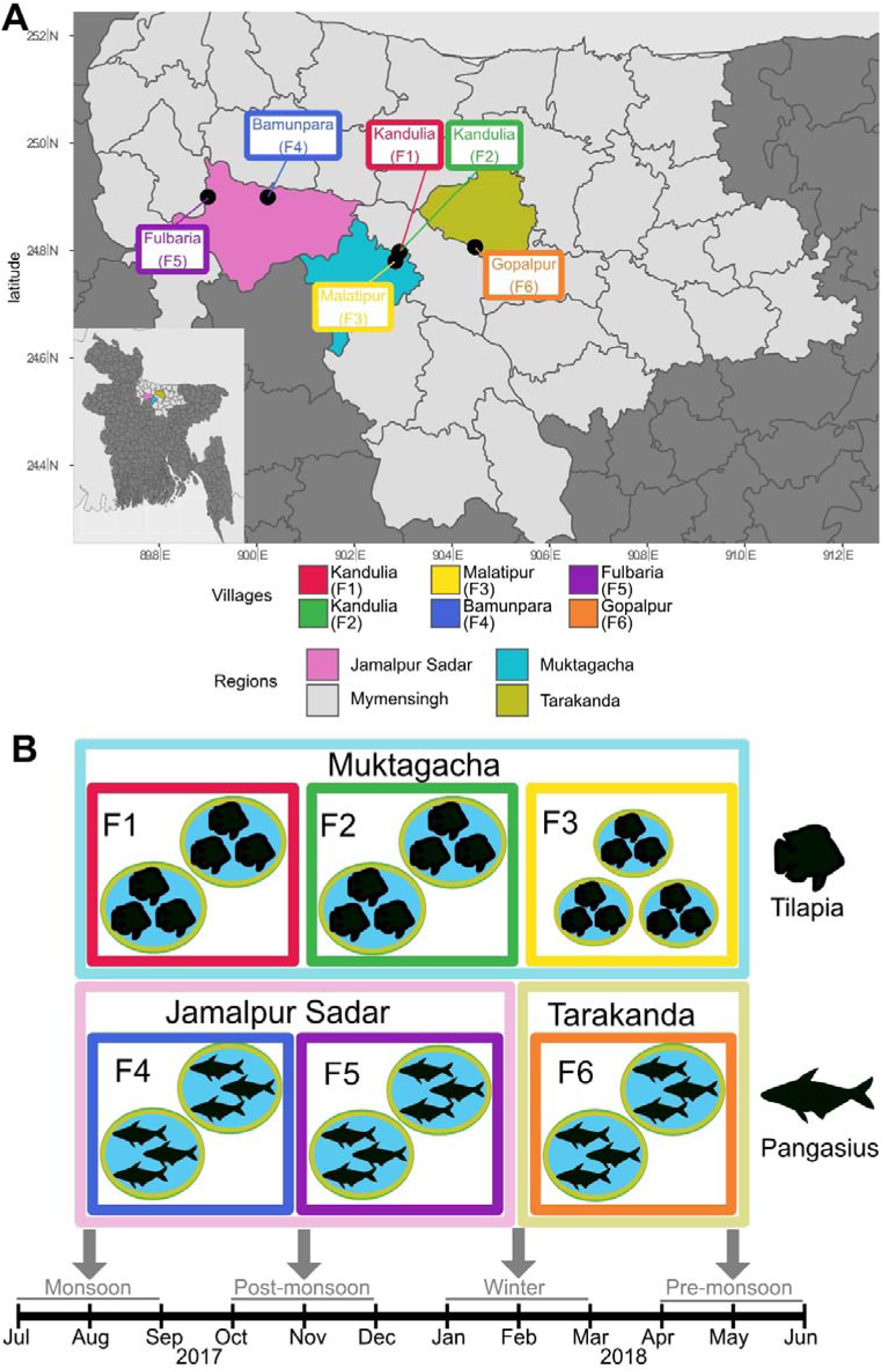
**A)** Regional map of the Mymensingh division of North-central Bangladesh. Fish farm locations within villages are labelled F1, F2, F3, F4, F5, F6 and corresponding Upazilas are highlighted with an inset showing a map of Bangladesh with the Mymensingh division highlighted in light grey. Farm locations are jittered slightly to preserve farmer anonymity. **B)** Sampling regime schematic with pond water samples collected and pooled from six different Bangladesh finfish aquaculture farms (two or three ponds per farm) in three Upazilas and over four seasonal periods.

### 2.2. Sample collection

Pond water samples were collected monthly between July 2017 and June 2018 from the same two or three ponds on each farm within the three Upazilas (geographical subdistricts) (Figure 1B). To collect microbial biomass, a maximum of 200 mL of pond water was passed through a filter (47 mm diameter, 0.4 μm pore size, 7060-4704, Whatman) using a syringe. In some cases, suspended particles clogged the filter before 200 mL of water passed through, resulting in a reduced amount of water filtered. Samples were collected in triplicate resulting in a maximum of 600 mL of filtered water per pond to obtain microbial biomass. To provide sufficient DNA for PCR-free shotgun metagenomic sequencing, samples collected across these monthly samplings were combined in the following manner: monsoon (corresponding to July to August 2017); post-monsoon (corresponding to October to November 2017); winter (corresponding to January to February 2018); pre-monsoon (corresponding to April to May 2018). This created a total of twenty-four samples consisting of six pooled pond water samples (1, 2, 3, 4, 5, 6) over four timepoints (Figure 1C). Filters were stored immediately in molecular grade ethanol and kept at ambient temperature until arrival in the UK where they were stored at -20°C until processing.

### 2.3. DNA extraction and sequencing

Ethanol was removed from filters by freeze-drying at -110°C (ScanVac CoolSafe Pro; FRE4578, SLS) and the filters transferred to storage at -80°C. DNA was extracted using an in-house CTAB/EDTA/chloroform method (Chaput, 2021; adapted from Bramwell et al., 1995; Lever et al., 2015). The full protocol is available at https://dx.doi.org/10.17504/protocols.io.bw8gphtw. DNA from pond water samples was cleaned using the Genomic DNA Clean & Concentrator kit (D4010, Zymo Research), quantified with the Qubit dsDNA BR Kit (Q32850, ThermoFisher Scientific), and submitted to the University of Exeter Sequencing Service for paired-end PCR-free shotgun metagenome sequencing on the NovaSeq 6000 with the S1 Reagent Kit (300 cycles). Raw sequencing reads were deposited in the European Nucleotide Archive (ENA) under the BioProject Accession Number PRJEB53918.

### 2.4. Bioinformatic assembly, classification and abundance

Scripts for all bioinformatic processes are available on GitHub (https://github.com/ash-bell/Bangladesh_ARG_analysis/). A publicly available bioinformatics pipeline from bbmap v38.90 (BBMap/pipelines/assemblyPipeline.sh) (Bushnell, 2013) was used to pre-process the shotgun metagenomic reads. Pre-processing involved adaptor sequences removal, read error correction and removal of human DNA. Reads were assembled using metaSPAdes v3.15.3 (Nurk et al., 2017) with the read normalisation step removed, as this step was not beneficial for our assembly. Predicted protein sequences were identified on contigs by Prodigal v2.6.3 in metagenomics mode (Hyatt et al., 2010). The Resistance Gene Identifier (RGI) v5.2.1 (Alcock et al., 2020) was used to identify ARGs from protein sequences using a custom amino acid database with in-house cut-offs determined by the Comprehensive Antibiotic Resistance Database (CARD) curators (Alcock et al., 2020). This was used in preference to nucleotide-based databases as protein translations have a more conserved homology. The gene adeF, an efflux pump, was removed from all analyses as an outlier as its presence was 24-fold higher than the mean of all ARGs within this dataset (not shown). This has been shown in a previous study where protein homology matches for adeF resulted in false positives and were removed from that analysis (Yao & Yiu, 2019). To measure contig, ARG and plasmid abundance, reads were mapped to their assemblies using BBMap (Bushnell, 2013) and normalised with the metric transcripts per kilobase million (TPM) calculated from CoverM v0.6.1 (Woodcroft & Newell, 2021). Mappings were only included if query sequences had over 70% coverage and reads mapped with at least 95% identity over 90% of a read’s length. Taxonomy for all reads and contigs was determined using Kaiju v.1.7.0 (Menzel et al., 2016) and Kraken2 v2.1.2 (Wood et al., 2019) using the NCBI non-redundant database (downloaded 11^th^ February 2022). Where read taxonomic identities conflicted between Kaiju and Kraken2, the lowest common ancestor was used. Translation from NCBI taxonomy identifiers to scientific names was performed using TaxonKit v0.8.0 (Shen & Ren, 2021). Reads were reassembled using metaplasmidSPAdes (Antipov et al., 2019) to look for contigs with overlapping ends, indicative of circular genomic regions, commonly viral or plasmid in origin. To remove potential viruses, putative plasmids identified as viral by VirSorter2 v2.2.3 (Guo, Bolduc, et al., 2021) and CheckV v0.8.1 (Nayfach et al., 2021) as outlined in their best practice protocol (Guo, Vik, et al., 2021) or identified as viral from Kaiju or Kraken2 were discarded. Putative non-viral plasmids were then checked against PlasClass v1st/Nov/2021 (Pellow et al., 2020) and any contig with a score of less than 0.5 was discarded as a false positive, as outlined in Pellow et al. (2020). Classification of gene function by the Kyoto Encyclopedia of Genes and Genomes (KEGGs) (Kanehisa & Goto, 2000) was performed using DRAM v1.3 (Shaffer et al., 2020) using default settings. Detection of single copy marker gene recombinase A (RecA) from metagenomic assemblies used a hidden Markov model (NCBI; TIGR02012.1) from the TIGRfam database (Haft et al., 2003) was performed using a 1e^-50^ cut-off via the tool hmmer v3.3.2 (Eddy, 2011).

### 2.5. Alpha Diversity

Relative abundance of reads at a genus level (with no measurement (NA) values replaced as 0) was used to determine alpha diversities of ponds using Shannon diversity from the R package vegan (Oksanen, 2022). A linear mixed-effects model from the Ime4 R package with Farm as the random effect was used to assess if alpha diversities were significantly different between seasons, with cultured fish type or location as the fixed effects.

### 2.6. Beta Diversity

All calculations were performed using the vegan package in R. Relative abundance of contigs at a genus level (with no measurement (NA) values replaced as 0) was used to calculate a Bray-Curtis distance matrix. A Nonmetric Multidimensional Scaling (NMDS) was used to visualise community taxonomic dissimilarity. Environmental factors, including geographical locations, main crop (fish species), season and date were fitted to the NMDS ordination plot with a goodness of fit test to determine how well environmental factors correlated with community structure. A PERMANOVA test was used to determine if the variance of pond water taxonomic compositions was significantly different when categorised by environmental factors. If a significant difference was observed, a pairwise PERMANOVA was performed to determine which environmental factor had a significantly different pond water taxonomic variance. This was performed through the R package pairwiseAdonis (https://github.com/pmartinezarbizu/pairwiseAdonis) where p-values were corrected for multiple re-testing using the Benjamini-Hochberg method.

### 2.7. Drug Class significance

To determine if resistance to any antibiotic drug class differed in abundance at any location or by crop type, a PERMANOVA test from the R package vegan was used. A pairwise PERMANOVA through the R package pairwiseAdonis was used to determine if any specific antibiotic drug class was significantly different in abundance from other sites. P-values for the pairwise PERMANOVA were corrected for multiple re-testing using the Benjamini-Hochberg method.

### 2.8. Pathogen search

To determine if a contig was from a putative pathogen, taxonomic identity at a genus level determined from Kaiju was searched against the American Biological Safety Association (ABSA) database of know pathogens to humans, animals and plants. Matches were defined as any contig having its genus level identity corresponding with known pathogens on the ABSA database.

## 3. Results

To investigate variation in the overall microbial composition in the fish ponds over seasons across the different Bangladeshi fish farm sites, we first analysed alpha and beta diversity metrics, and community compositions.

### 3.1.1. Relative microbial taxonomic abundance in the pond waters

A mean of 47.37% of all reads across all samples could not be classified at any taxonomic level due to their absence (of similar sequences) in the NCBI’s nr database. Overall, taxonomically identified reads were comprised of Bacteria (94.65%), Viruses (2.41%), Eukaryota (2.14%) and Archaea (0.80%). The identified bacterial reads were dominated by Proteobacteria (30.8%), followed by Planctomycetes (18.9%) and Actinobacteria (15.4%) phyla (Figure 2). Of all identified Archaeal reads, the most abundant Archaea phyla were the Euryarchaeota (50.5%) (Supplementary Figure 1). Within the viral reads, Uroviricota were the most common viruses (72.6%) (Supplementary Figure 2). Almost all Eukaryotic reads were classified as human with a small number of reads from other Eukaryotic species (<0.000001% relative abundance). No differences were found between ponds at the domain (Bacteria, Archaea, Eukaryota and Viruses) level when assessing the phylum composition of each pond, indicating all samples had similar basic community structures.

**Figure 2.**
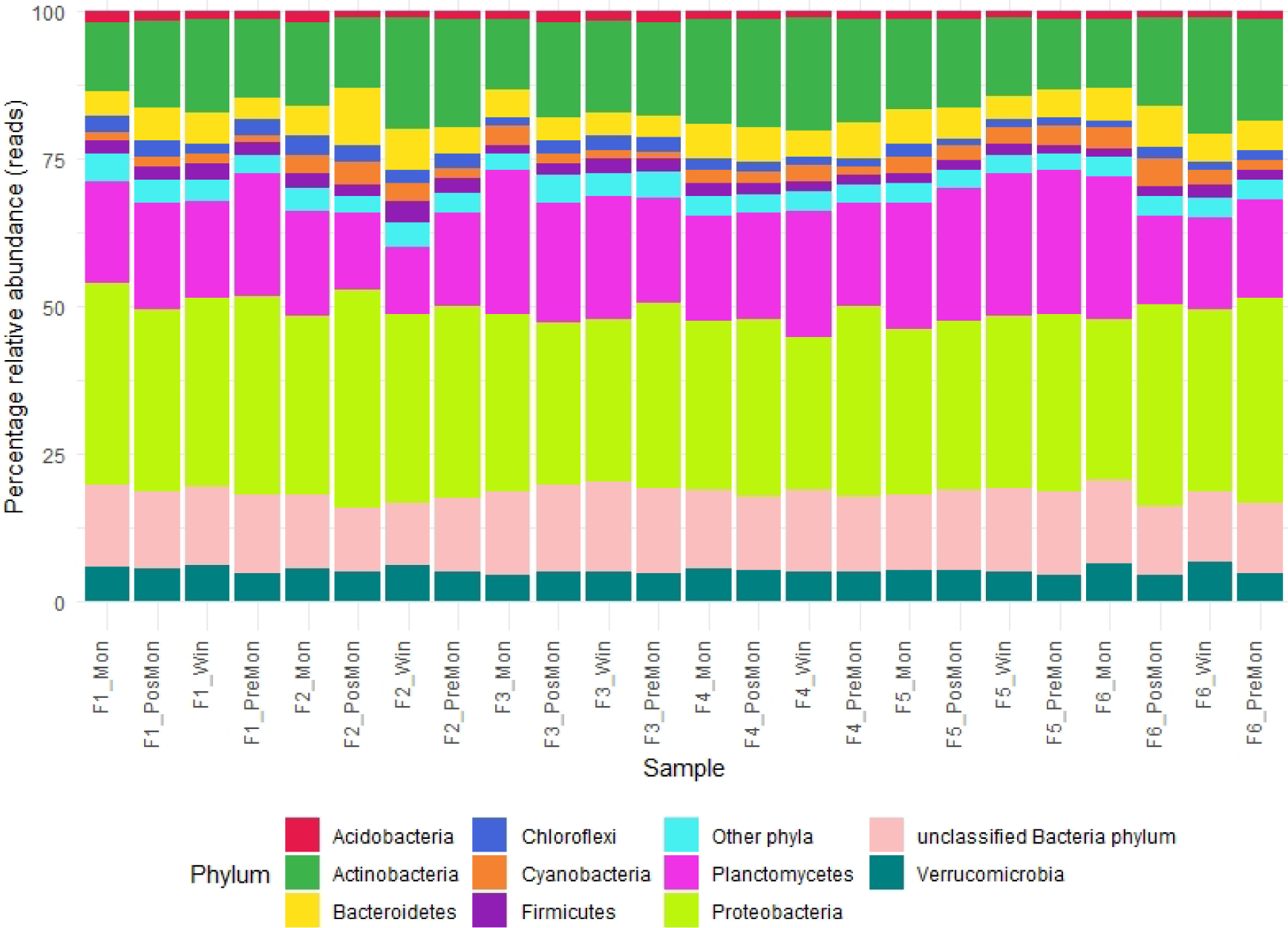
Relative abundance plot of taxonomically classified reads of bacteria in samples collected from six different Bangladesh finfish aquaculture farms in three Upazilas.

### 3.1.2. Pond microbial alpha diversity

Microbiological alpha diversity measured using Shannon diversity did not differ between different farms, main crop type (fish species), village, Upazila or season (Figure 3) as determined using a linear (mixed) model (Supplementary Table 1).

**Figure 3.**
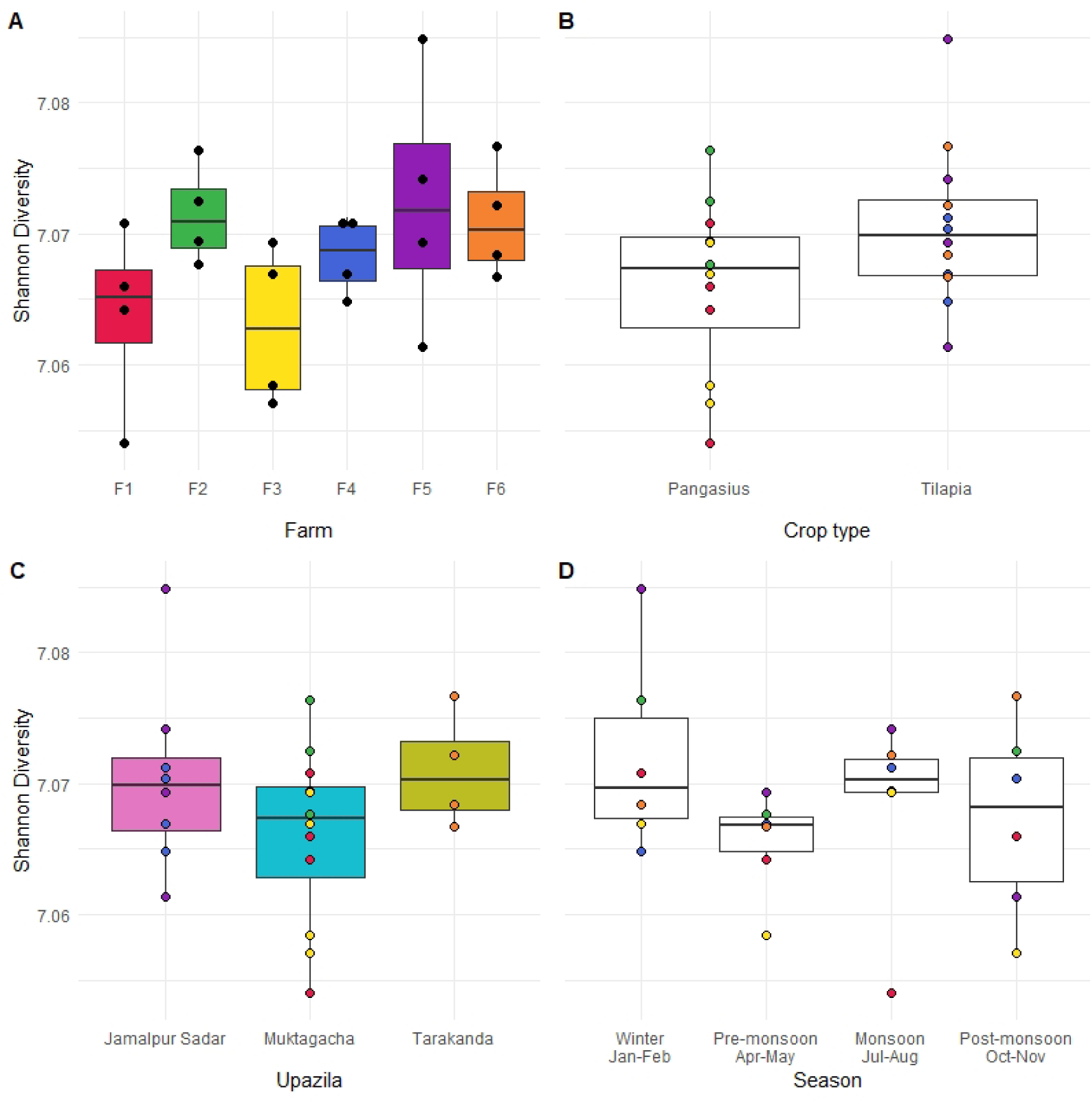
Microbial Shannon diversity for Bangladeshi finfish aquaculture pond waters grouped according to **A)** Farm location **B)** Fish species **C)** Upazila **D)** Season, illustrating no significant differences between ponds. Colours are linked with Figure 1.

### 3.1.3. Upazila correlates with differing pond microbiome community structures

Nonmetric Multidimensional Scaling (NMDS) was used as a dimension reduction technique to visualise differences in microbial community compositions as a single point with colours indicating samples from the same farm or Upazila. Pond microbial beta diversity at the genus level did not cluster together for any one farm location indicating pond ‘replicates’ were as dissimilar in their microbial composition as farms from a different location (PERMANOVA, R^2^ = 0.266, p = 0.11) (goodness of fit of environmental factors (coloured arrows), R^2^ = 0.314, p=0.185) (Figure 4A). Associations between beta diversity at farms and environment factors (tilapia/pangasius main crop, village, Upazila and season) found only Upazila significantly correlated (but weakly so, Figure 4B) (PERMANOVA R^2^ = 0.147, p=0.024). Pairwise PERMANOVA tests indicated significant differences in the beta diversity between Muktagacha compared with Tarakanda only (pairwise PERMANOVA R^2^ = 0.162, p= 0.027).

**Figure 4.**
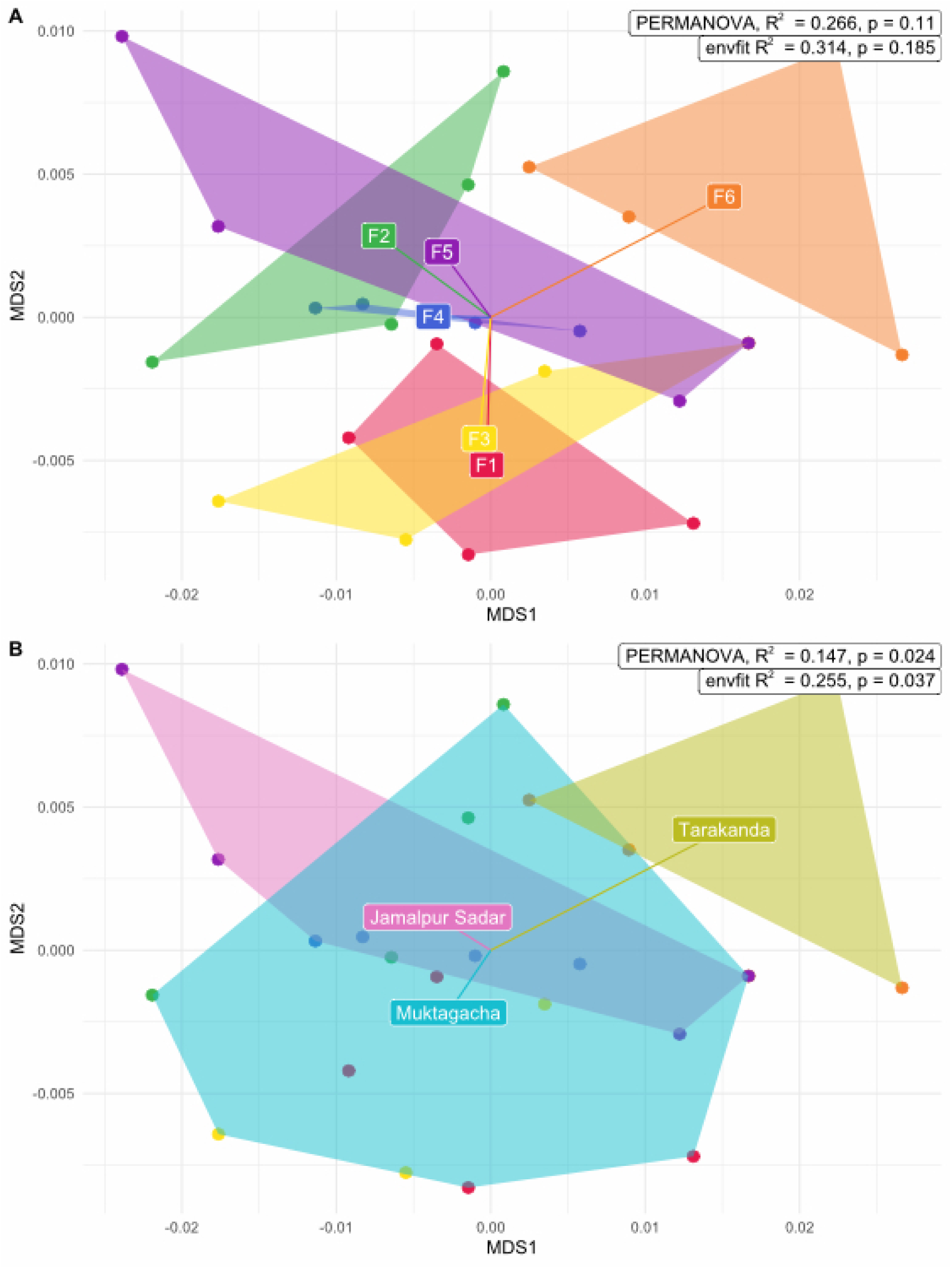
Nonmetric Multidimensional Scaling (NMDS) plot of beta diversity (genus level) of fish aquaculture ponds. **A)** Farm sites with lines from the centre of the plot to centroids depicting the direction of grouping variables (farms). The overall taxonomic composition did not differ between the different farm sites. **B)** Upazilas with lines from the centre of the plot to centroids depicting the direction of grouping variables (Upazila). Upazilas microbial pond compositions are significantly different from one other. Colours are linked with Figure 1.

### 3.1.4. Pond microbiome functional diversity

Analysis of the functional profile of genes within each sample using KEGGs did not reveal a significant difference by PERMANOVA between season, Upazila or fish type (Figure 5) in line with findings that microbial community compositions were not significantly different from each other (Figures 3 and 4).

**Figure 5.**
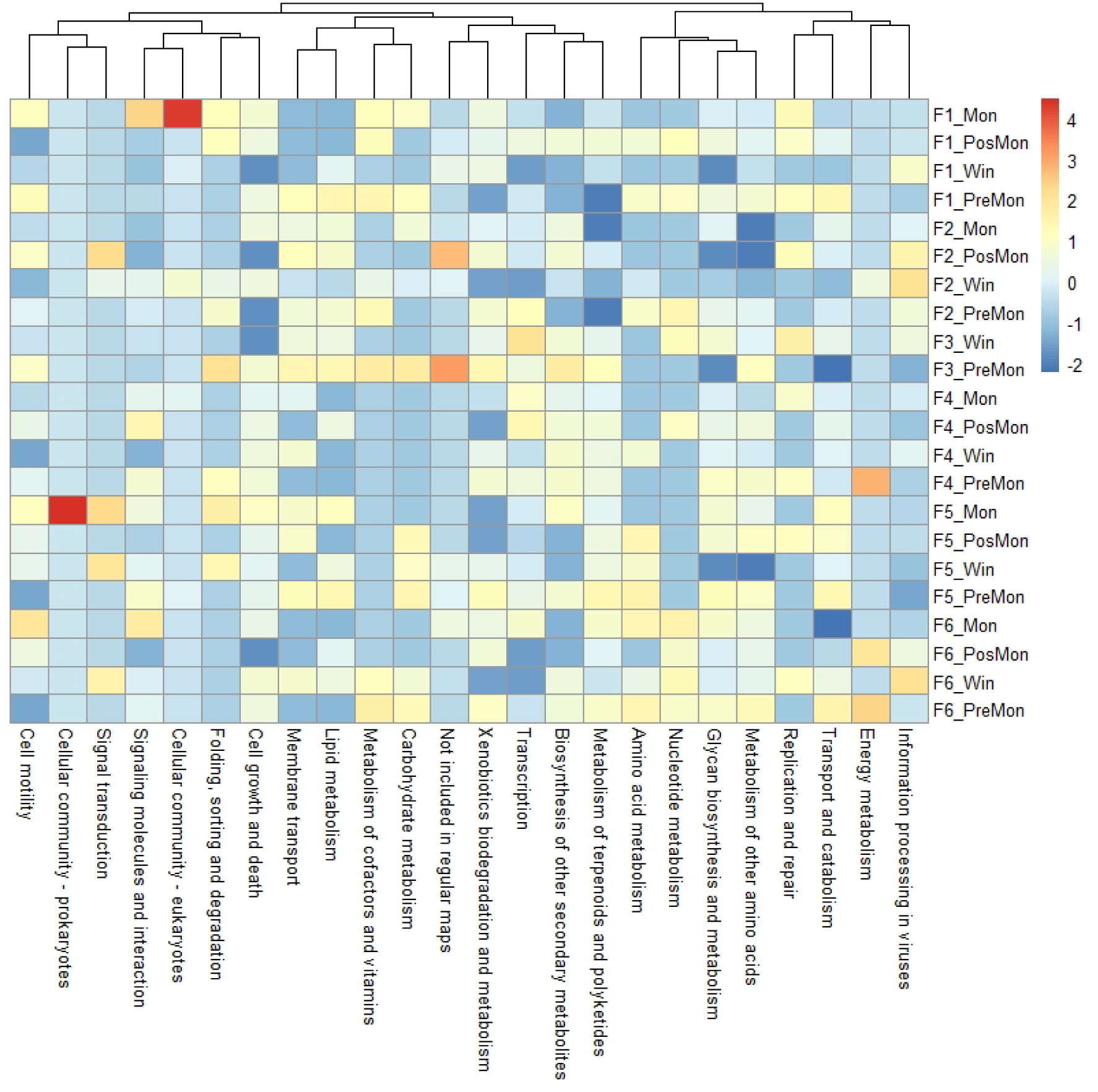
The abundance of genes was classified using KEGGs by sample within Bangladeshi finfish aquaculture ponds for samples collected from six different Bangladesh finfish aquaculture farms over four seasonal periods. Fish farm locations are labelled F1, F2, F3, F4, F5 and F6. Seasons are labelled monsoon (corresponding to July to August 2017); post-monsoon (corresponding to October to November 2017); winter (corresponding to January to February 2018); pre-monsoon (corresponding to April to May 2018). The abundance of KEGGs is summarised by the metabolic process they belong to using the metric transcripts per kilobase million (TPM). Values are centred by subtracting sample means and scaled by dividing by sample standard deviation (3 to -1).

### 3.2.1. Multiple ARGs conferring resistance to various antibiotics

The abundance and type of ARGs found within Bangladeshi fish pond water samples were identified and matched against a curated set of ARGs. This was done using a bioinformatically tested cut-off as determined by the Resistant Gene Identifier (Alcock et al., 2020). Matched ARGs were grouped by the antibiotic class(es) against which they provide resistance. Overall, there were ARGs resistant to 18 classes of antibiotics (out of 36 possible classes; CARD database, (Alcock et al., 2020)). Resistance to aminoglycosides was the most abundant drug class and along with sulphonamides, which was found on all six farms (Supplementary Figure 3, Supplementary Table 2). As there was a weak but significant difference in microbial community composition when grouped by Upazila, we looked to see if the same was true for ARG abundance (Figure 6).

**Figure 6.**
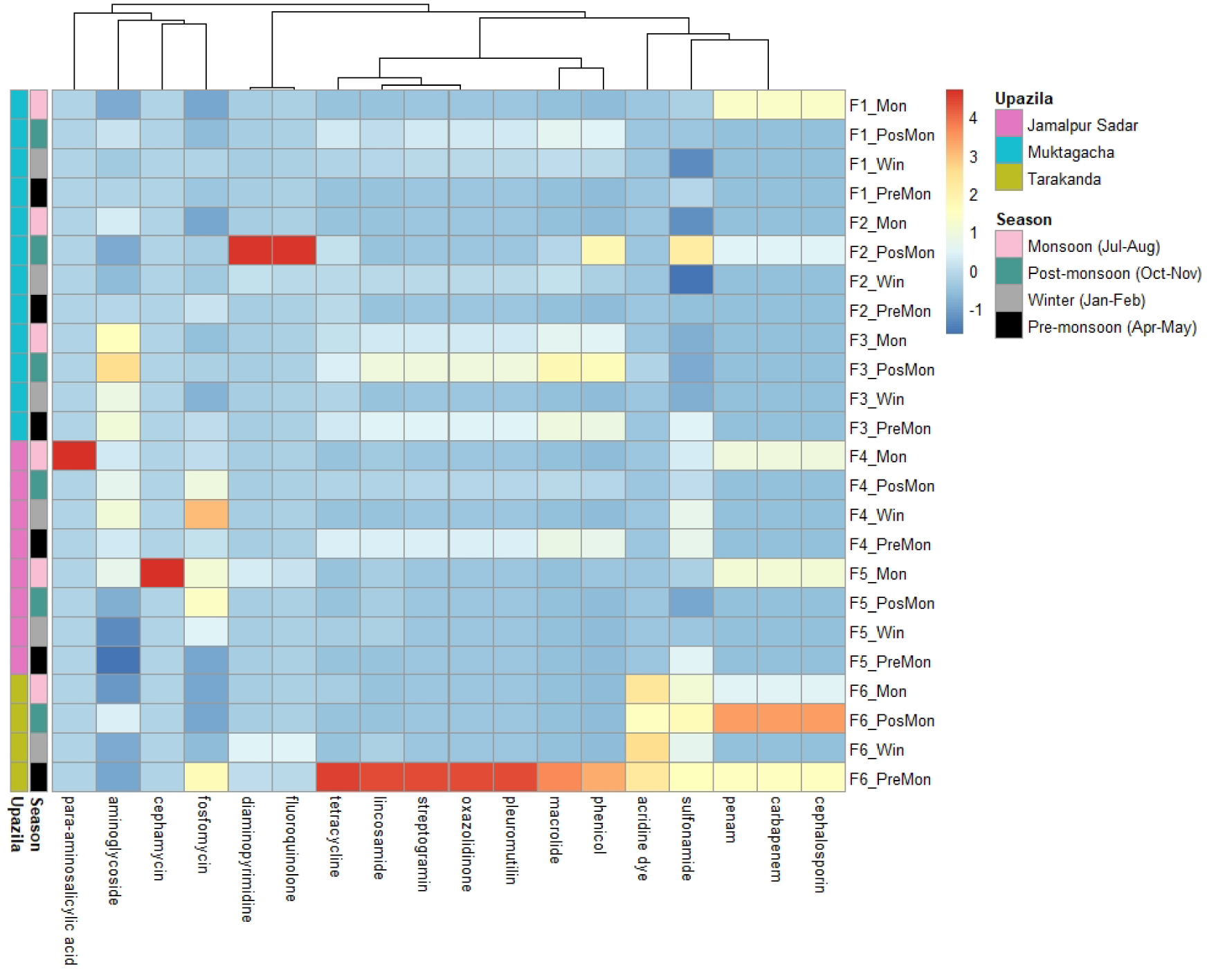
The abundance of ARGs to resistance against specific antibiotic drug classes within Bangladeshi finfish aquaculture ponds for samples collected from six different Bangladesh finfish aquaculture farms in three Upazilas and over four seasonal periods. The abundance of ARGs is summarised as the class of antibiotics they confer resistance to using the metric transcripts per kilobase million (TPM). Values are centred by subtracting sample means and scaled by dividing by sample standard deviation (4 to -1).

Investigating the abundance of specific ARGs, the rspL gene encoding ribosomal protein S12 with mutations resulting in resistance to aminoglycosides (e.g., streptomycin) was the most abundant gene (found in 23 of 24 samples). The rpsL gene is a universal single-copy marker gene and its high abundance may be an artefact of its ubiquitous nature. Therefore, its abundance was compared to the recA single copy marker gene to provide a baseline of expected abundance for single marker copy genes within each metagenome. RecA had an average abundance of 319.098 TPM (Supplementary Table 3), 46.44807-fold higher than the mean mutated rpsL abundance (6.87 TPM) conferring resistance to aminoglycosides. This suggests that read mapping and ARG identity tools are accurate in their assessment of ARG abundance and identity. The gene sul1 occurred in all ponds and sul2 was found in twenty-two of the twenty-four samples; both genes confer resistance to sulphonamides. Gene murA, conferring resistance to fosfomycin, was also notably abundant, identified in 19 of the 24 samples. Gene poxtA, implicated as a multi-drug resistant gene, was a further abundant gene present in nine of the twenty-four samples (Supplementary Figure 4, Supplementary Table 4). A PERMANOVA showed resistance to antimicrobial drug class or specific genes was not significantly enriched in any location, crop type or season. A similar antibiotic resistant gene compositional profile was present in all ponds at all locations and timepoints (Figure 6).

### 3.2.2. Multiple ARGs found on contigs associated with pathogenic organisms

Screening for ARGs encoded by pathogens using the ABSA database (for known human, animal and plant pathogens) identified 16 putative pathogens encoding resistance to 12 different antibiotic drug classes. All pathogens detected are known opportunistic or pathogenic to humans. Fosfomycin resistance was found in Mycobacterium spp., sulphonamide resistance in Escherichia coli and Salmonella enterica, aminoglycoside resistance in Klebsiella pneumoniae. Pseudomonas spp. encoding multiple resistances were found at two farms. ARGs AAC(6’)lb10 and CAM-1 were found on Pseudomonas spp. on farm five conferring resistance to aminoglycosides, carbapenems, cephalosporins, cephamycins and penams. On farm two, ARG rsmA was found on Pseudomonas spp. conferring resistance to fluoroquinolones, diaminopyrimidines and phenicols (Figure 7).

**Figure 7.**
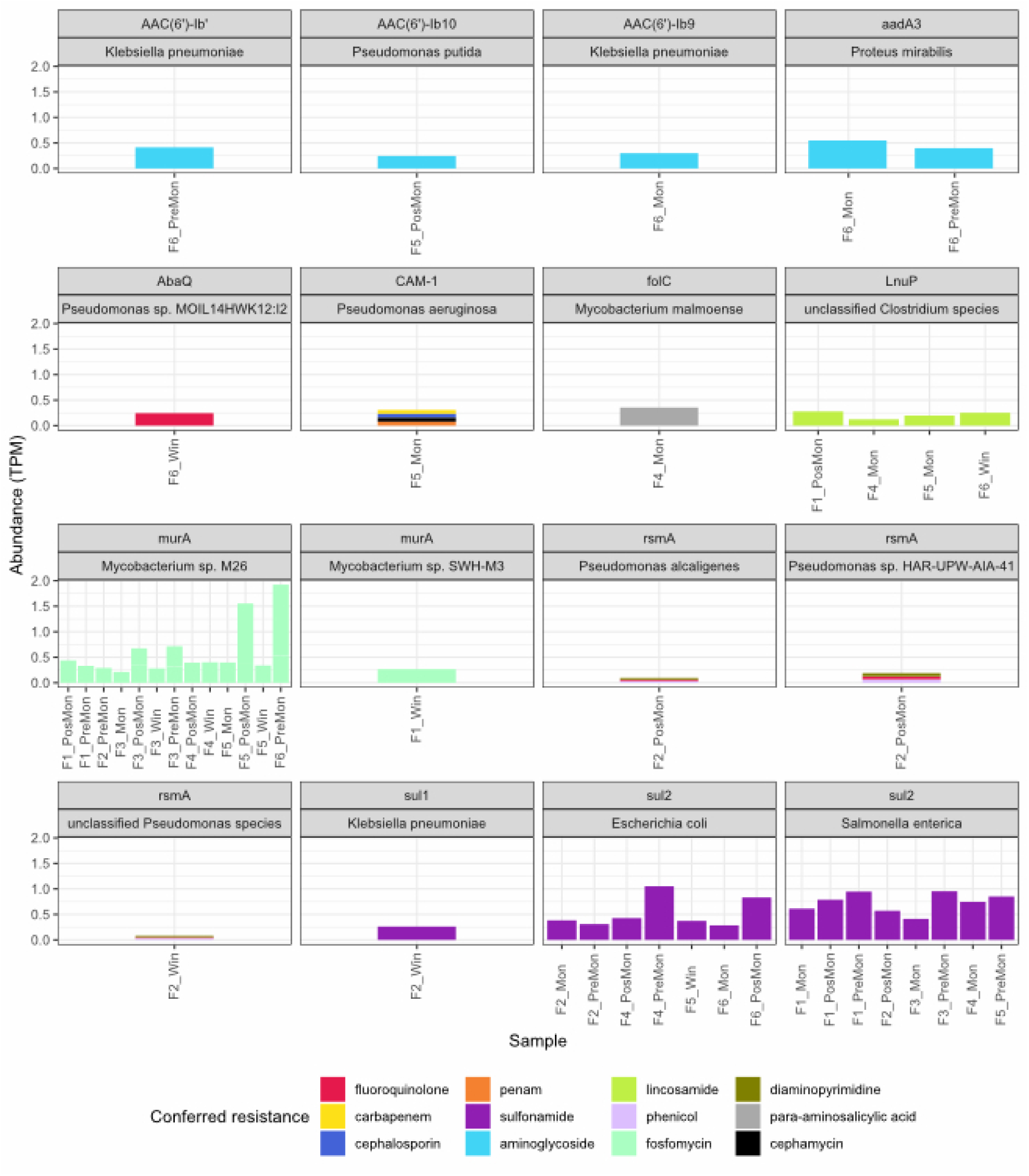
Abundance of ARGs conferring resistance to specific antibiotic drug classes summarised by ARG and species.

### 3.2.3. ARGs conferring resistance to sulphonamides and lincosamides on mobile genetic elements

A total of 6986 non-viral circular contigs were identified as putative plasmids through PlasClass. A total of five plasmids encoding ARGs lnuC, sul1 and sul2 were identified conferring resistance to lincosamides and sulphonamides antibiotics respectively (Supplementary Figure 5). Multiple genes encoding with mobile elements (Intl1, transposases) were found on these circular contigs.

#### Bacterial genomes

ARGs were not encoded universally or equally across all bacterial phyla. Aminoglycosides were almost exclusively encoded by the Phylum Actinobacteria and included multiple ARGs with multiple drug resistances. Planctomyces, Proteobacteria and Verrucomicrobia phyla were more likely to encode for tetracycline and fluoroquinolone resistance exclusively (Supplementary Figure 6).

## 4. Discussion

By applying metagenomic techniques we show microbial profiles of Bangladeshi fish aquaculture ponds across six farms in the Mymensingh division had a similar diversity. Microbiomes were dominated by Proteobacteria for bacteria, Euryarchaeota for archaea and Uroviricota for viruses, with no obvious seasonal differences. Proteobacteria were also the most common bacterial phylum found in aquaculture earthen finfish ponds in Malawi (McMurtrie et al., 2022) and China (Liu et al., 2020; Qin et al., 2016). In another study characterising pond microbiomes across districts in Bangladesh (Mymensingh, Shariatpur, and Dhaka), Cyanobacteria were the most common phylum, however, Proteobacteria were the most highly abundant bacteria phyla in five of seven water samples in the Dhaka district (McInnes et al., 2021). Of all environmental factors assessed in our study (fish species farmed in the ponds, pond location, and season/sampling timepoint), only Upazila correlated with microbial diversity. This supports previous work indicating that geography influences the microbiota composition of aquaculture ponds in Bangladesh (McInnes et al., 2021) and Malawi (McMurtrie et al., 2022). This is also true in freshwater lakes (Yang et al., 2019) and open oceans (Sehnal et al., 2021) where geography is correlated with aquatic microbial diversity.

In our study, we found that larger geographical features such as districts and Upazilas have a stronger bearing on pond microbiological compositional profiles compared to smaller geographical scales (i.e., neighbouring villages). The type of fish crop (tilapia vs pangasius) had little influence on the pond microbiota composition, indicating that these fish species do not play a major role in affecting the microbiota in these aquaculture pond systems. Looking at the physiochemistry of tilapia and pangasius fish ponds across multiple Bangladeshi districts, significant differences were detected in dissolved oxygen and nitrate (but not pH, orthophosphate or ammonia) levels between tilapia and pangasius farms (Islam et al., 2021). Although an apparent higher level of E. coli and total coliform abundance was indicated for tilapia (versus pangasius) ponds, this was not statistically significant (Islam et al., 2021). This correlates with our findings that fish pond microbiological community composition is not significantly influenced by the fish species they contained. Another study assessed the microbial composition of pond water from the cultivation of five different fish species (grass carp (Ctenopharyngodon Idella), adult largemouth black bass (Micropterus salmoides), juvenile largemouth black bass, hybrid snakehead (Channa spp.) and tilapia (Oreochromis niloticus)), found only in a minority of cases (grass carp versus adult largemouth black bass; grass carp versus hybrid snakehead) differences in alpha diversity (Liu et al., 2020). In addition, no differences were seen in bacterial community composition as measured by relative abundance or in PCA plots depicting beta diversity suggesting again, that overall fish species was not a driving factor in the microbial diversity of pond water (Liu et al., 2020).

Interestingly, we did not detect any seasonal variation in microbial diversity, including during the monsoon season. Similarly, Azmuda et al. (2019) looking at the microbial community composition of Bangladesh ponds and lakes across different seasons, found no variation in microbiomes despite seasonal changes in physicochemical factors (temperature, pH, salinity, conductivity). This finding is somewhat surprising considering bacteria in water bodies are sensitive to changes in physicochemical factors, such as pH and temperature, which are correlated with seasonality (Gilbert et al., 2011; Newton et al., 2011; Smyth et al., 2014). We suggest that the pond aquaculture practices themselves, including the feeding and probiotic treatment regimes, and the use of lime and other chemicals to maintain physicochemical factors suitable for supporting the growth of cyanobacteria (a feed source for cultured fish, such as tilapia) (Jahan et al., 2015) are more influential in determining pond water bacterial community compositions than season or the fish species stocked.

Using metagenomic techniques we found high ARG abundance and diversity within Bangladeshi fish ponds conferring resistance to eighteen different antibiotic classes. ARGs within these fish ponds were present in putative plasmids, of which ARGs encoding resistance to twelve different antibiotic drug classes, were found within microbial taxa that include human and animal pathogens. The most abundant antibiotic drug resistance class was aminoglycosides (as well as sulphonamides) and found to occur across all the farm sites analysed. These findings correspond with a study by McInnes et al. (2021) where ARGs encoding resistance to sulphonamides, macrolides, and aminoglycosides were the most common antibiotic resistance classes found in rural Bangladeshi surface waters, including ponds. Overall, although there was a weak correlation between microbial composition and location (Upazila), the ARG content was similar in all ponds sampled within the Mymensingh Division of Bangladesh. As for the microbial composition, no significant seasonal differences in ARG compositions were seen in the Bangladeshi fish ponds or between the ponds culturing the different fish species (pangasius or tilapia). ARGs were not encoded universally or equally across all bacterial phyla, in line with findings from the human gut microbiome that there are increased barriers for mobile genetic element transfers as phylogenomic distances increase (Forster et al., 2022; Jiang et al., 2019).

ARGs conferring resistance to eight different drug classes over the entire dataset were found in the genus Pseudomonas with five classes in one farm alone. Pseudomonas spp. include opportunistic human pathogens such as P. aeruginosa, the fish pathogen P. fluorescens and the plant pathogen P. syringae. ARGs found in Pseudomonas spp. included CAM-1 (conferring resistance to beta-lactams including cephalosporins, carbapenems, cephamycins and penams), AAC(6’)-Ib-10 (conferring resistance to aminoglycosides) on farm five and rsmA (conferring resistance to fluoroquinolones, phenicols and diaminopyrimidines) on farm two. Pseudomonas infections are normally cleared with antibiotics, such as fluoroquinolones (e.g. ciprofloxacin) (Poole, 2011) or aminoglycosides (e.g. kanamycin) (Poole, 2004), with carbapenems used as a last resort, due Pseudomonas’s innate resistance to other types of antibiotic classes, such as macrolides and tetracyclines (Morita et al., 2013). Within this dataset, we show that Pseudomonas spp. has ARGs conferring resistance to all three commonly used antibiotic classes. Bacterial acquisition of ARGs conferring multi-drug resistance can reduce antibiotic effectiveness, poses risks to effectively treating infections and increase human and crop mortality.

Our study shows that ARGs differ more in their compositional profile across larger geographical districts and Upazilas compared to smaller geographical features such as neighbouring villages. Similarly, ARG profiles were recently shown to be significantly different between rural and urban fish farms from different districts within Bangladesh but not within the same district (Lassen et al., 2021). Their study also highlighted a north-south division in the use of farm-made vs commercial feed and postulated that these farming practices were responsible for the different ARG profiles in northern versus southern districts (Lassen et al., 2021). As farmers within our study used both commercial sinking and floating feed irrespective of fish species cultivated, we cannot support Lassen et al.’s (2021) claim.

In summary, we find a high diversity of ARGs present in all finfish ponds sampled in the Mymensingh division of Bangladesh, with the most common being aminoglycosides and sulphonamides resistance genes. ARGs were furthermore identified in putative mobile genetic elements. ARG compositions and abundances within these fish ponds did not differ over season or with crop type (fish species). There was a weak but significant difference in pond water compositions across the larger geographical classifications (Upazila vs village). Given the limited geographical difference and lack of seasonal differences in the microbiota and ARG profiles, we suggest that these are driven by farming practices. These traditional earthen fish ponds are filled predominantly using rainwater (but receive groundwater in dry conditions), receive similar chemical treatments and foodstuffs (often laced with antibiotics) and are often heavily stocked with fish. Similar farming practices may create similar water physicochemical (typically, eutrophic with low and diurnal fluctuating oxygen and pH) in turn likely creating similar microbial selection pressures and thus homogenising microbial diversity. At the start of the pre-monsoon season (March/April), when fish are restocked, fish pond water is often prophylactically treated with antibiotics (Jahan et al., 2015). Combined with antibiotically laced food administered during multiple annual restocking events, this can reduce microbial diversity and select for AMR bacteria (Manyi-Loh et al., 2018; Murray et al., 2019). More studies are needed, however, to assess to what extent antibiotic usage is directly correlated with increased ARG abundance and reduced antibiotic efficacy in fish farms in Bangladesh. The current study highlights the need for a more informed approach to antibiotic use in the earthen pond aquaculture systems in Bangladesh (and globally) to help minimise ARG development, especially given the drives to intensify production with the inevitable increase in pathogen prevalence and disease outbreak likelihood.

## 5. Supplementary data

**Supplementary Figure 1.**
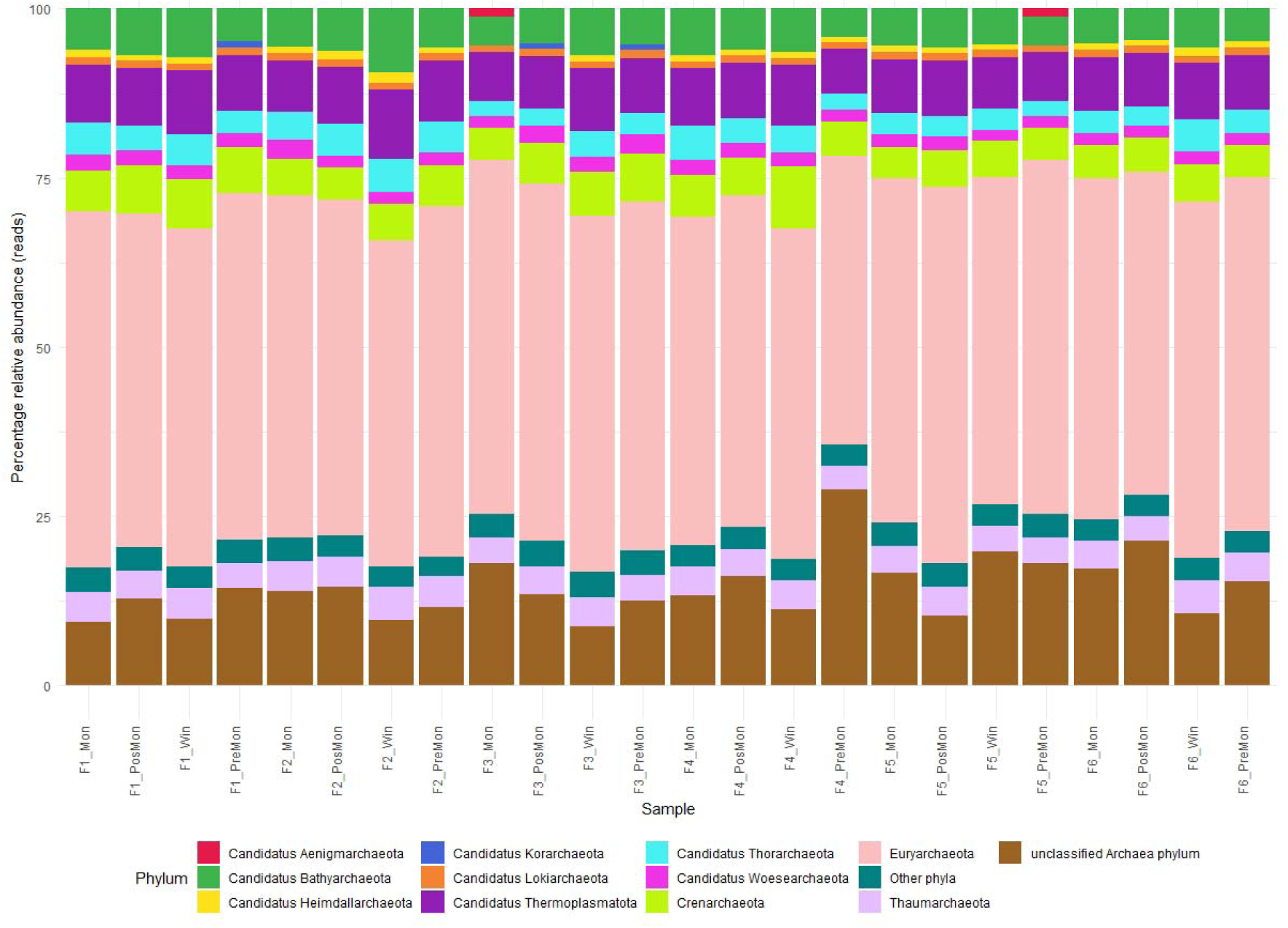
Relative abundance plot of taxonomically classified reads of Archaea in samples collected from six different Bangladesh finfish aquaculture farms in three Upazilas.

**Supplementary Figure 2.**
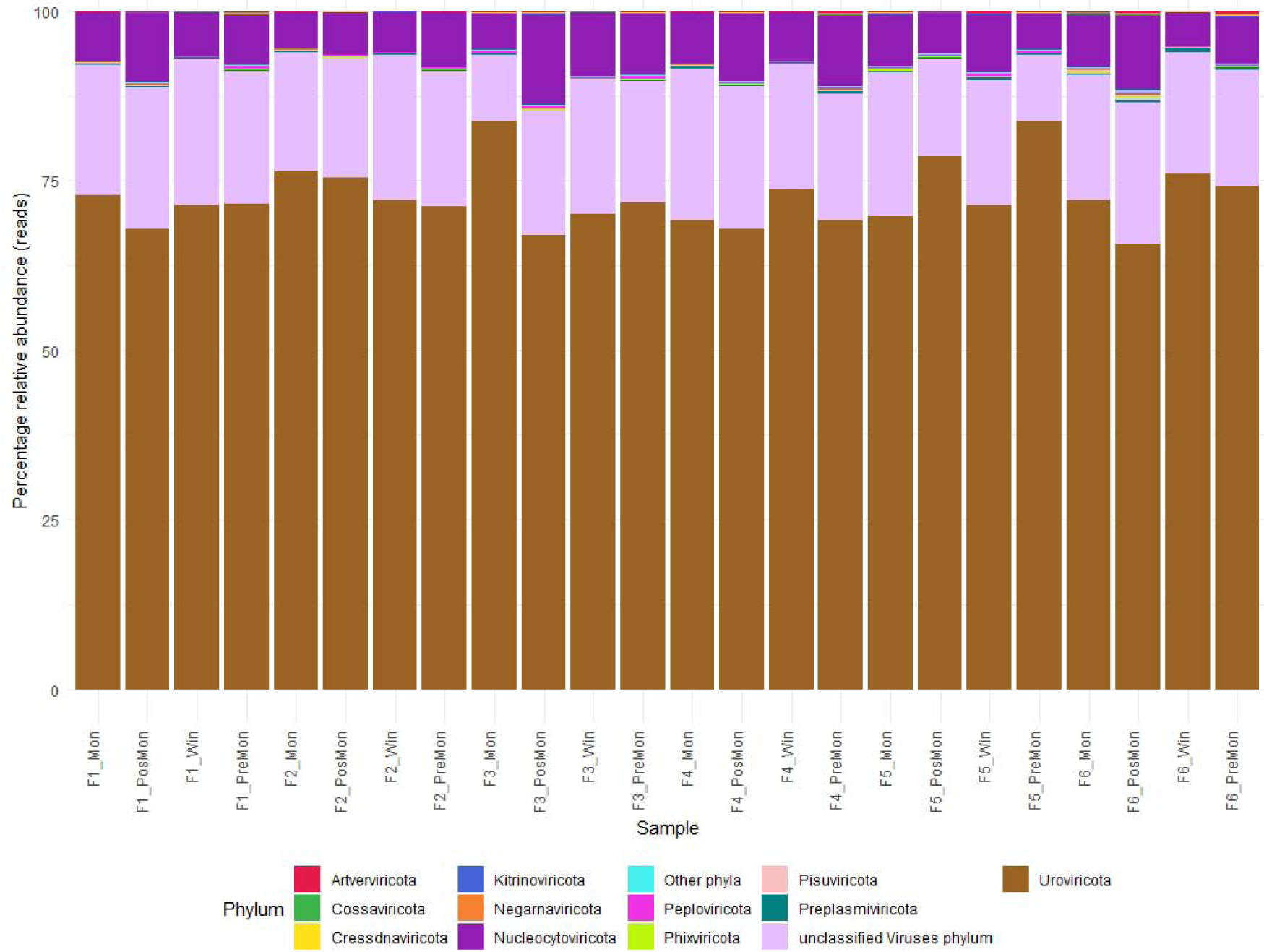
Relative abundance plot of taxonomically classified reads of viruses in samples collected from six different Bangladesh finfish aquaculture farms in three Upazilas.

**Supplementary Figure 3.**
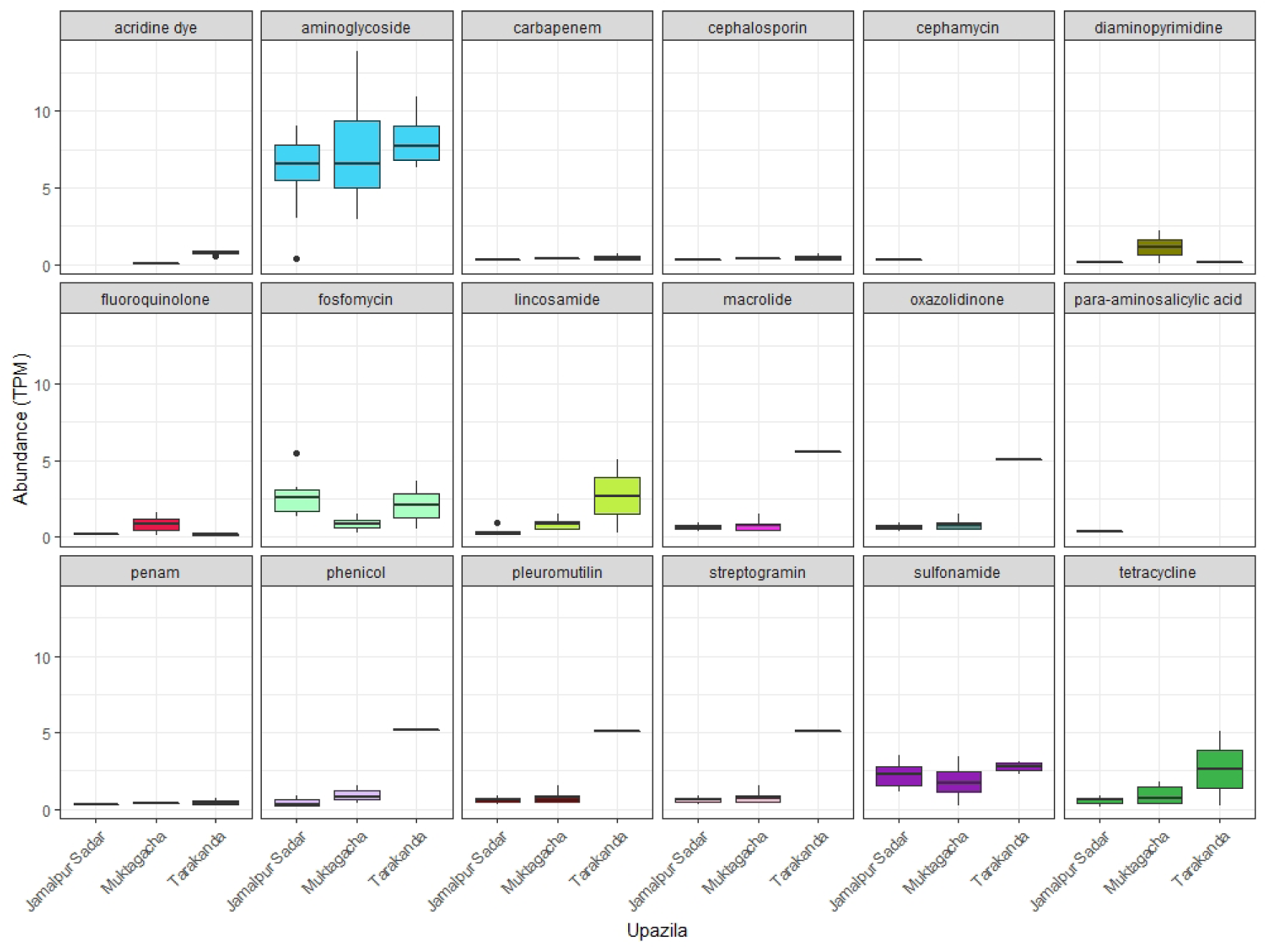
Abundance of ARGs found within six different Bangladesh finfish aquaculture farms in three Upazilas. Abundance of ARGs is summarised by antibiotic drug class they confer resistance to and grouped by Upazila they were found in. ARGs abundance metric is Transcripts per kilobase Million (TPM).

**Supplementary Figure 4.**
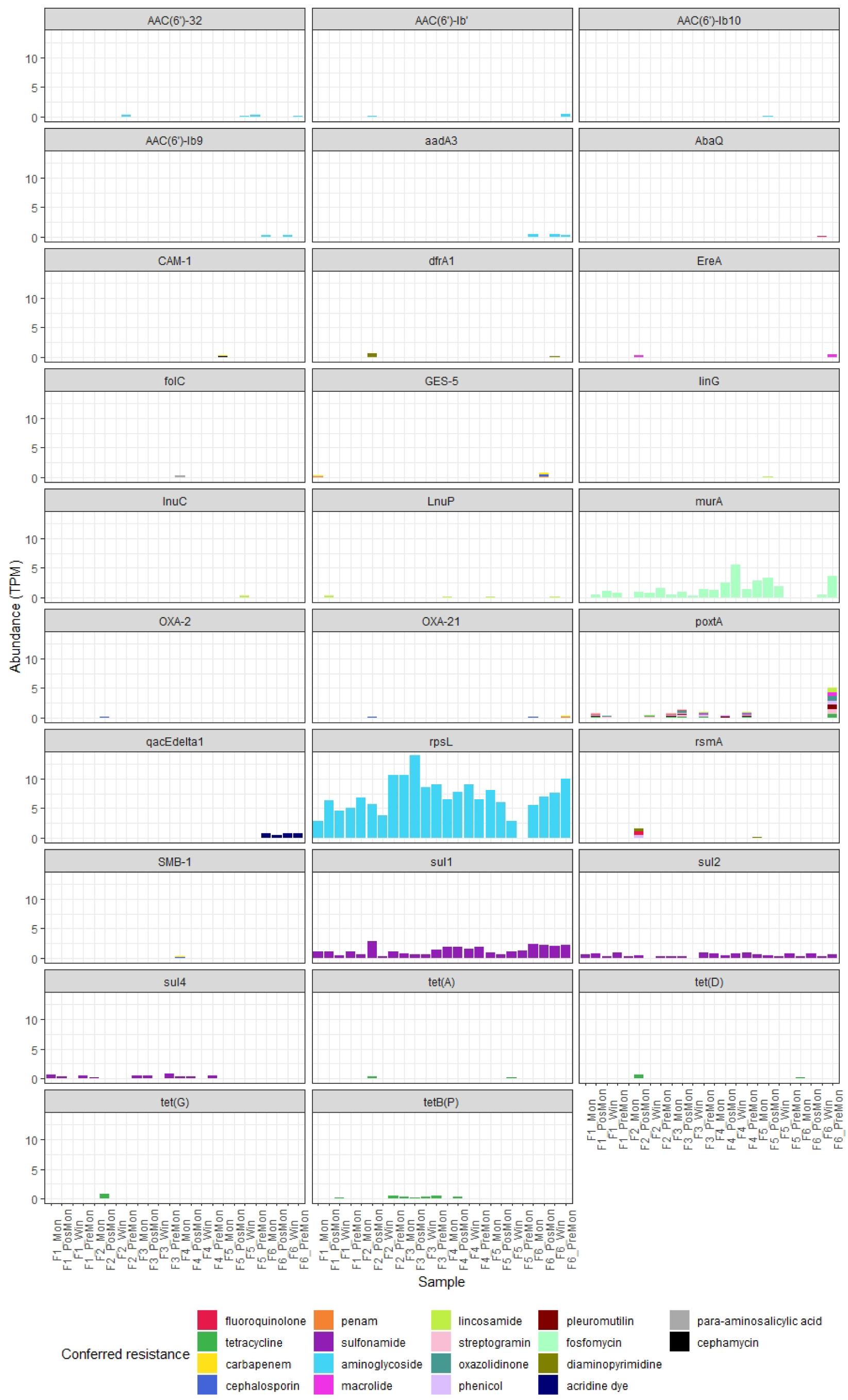
Abundance of ARGs found within six different Bangladesh finfish aquaculture farms in three Upazilas and over four seasonal periods. ARGs abundance metric is Transcripts per kilobase Million (TPM).

**Supplementary Figure 5.**
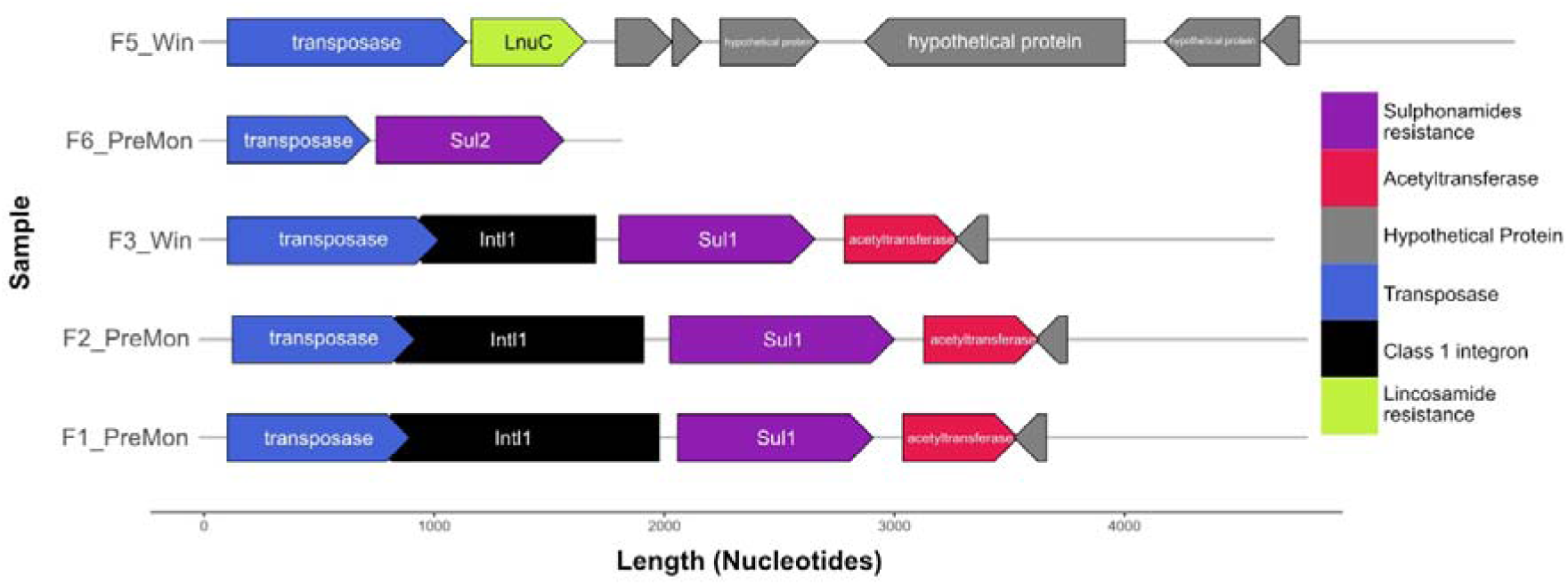
Gene annotation of five recovered plasmids with mobile genetic elements and antimicrobial resistant genes conveying resistance to antibiotic drug classes.

**Supplementary Figure 6.**
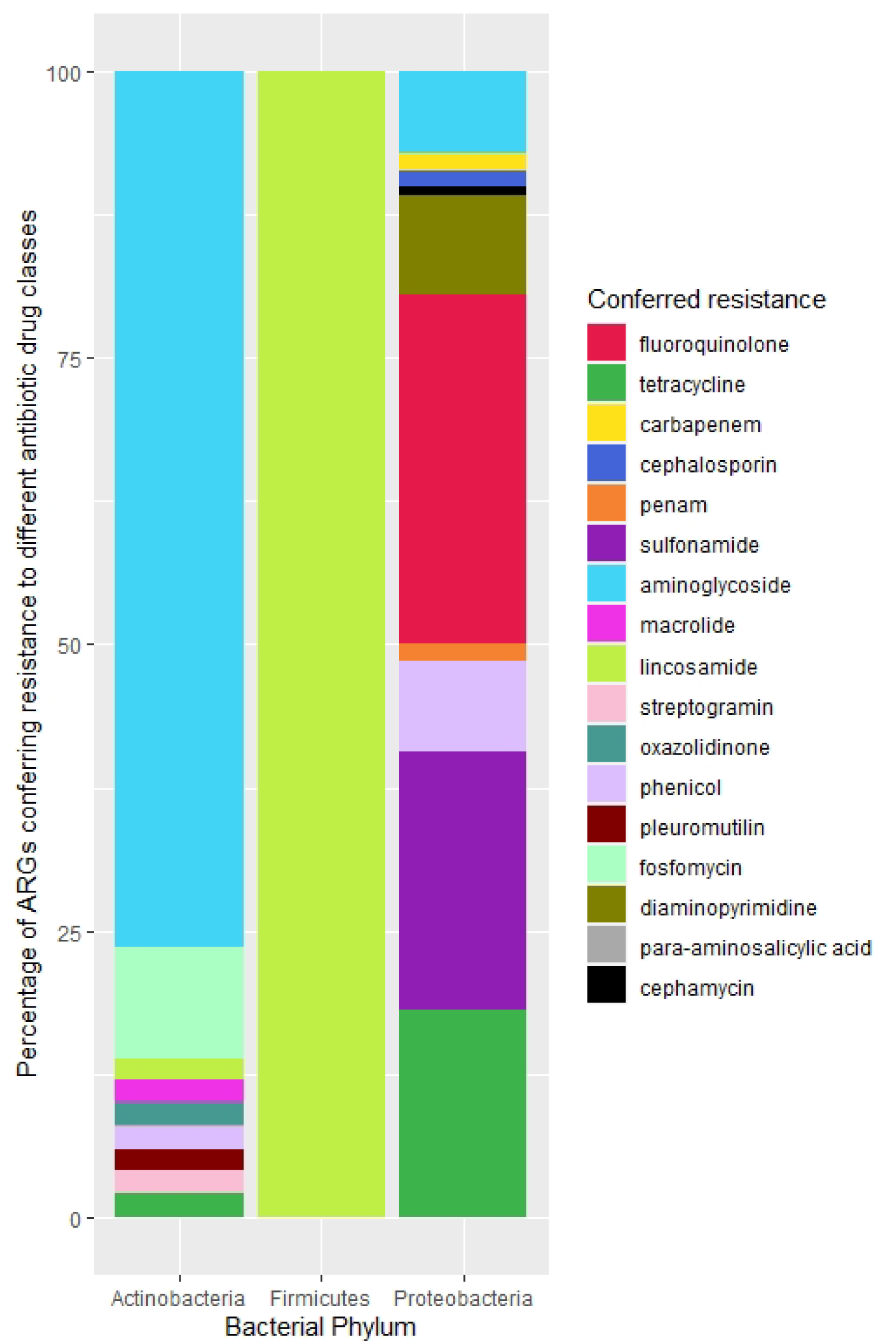
Percentage abundance of ARGs encoding resistance to specific drug classes summarised by bacterial phylum.

**Supplementary Table 1:** Linear mixed model results for Shannon alpha diversity comparing different environmental metrics

| Metric | Model | Random effect | Random Effects Standard Deviation | Fixed Effect | Beta | Confidence interval 2.5% | Confidence interval 97.5% | p value |
| --- | --- | --- | --- | --- | --- | --- | --- | --- |
| Shannon Diversity | Linear model | N/A | N/A | FarmerF1 (Intercept) | 7.06 | 7.06 | 7.07 | 0 |
|  |  |  |  | FarmerF2 | 0.01 | 0 | 0.02 | 0.09 |
|  |  |  |  | FarmerF3 | 0 | -0.01 | 0.01 | 0.85 |
|  |  |  |  | FarmerF4 | 0 | 0 | 0.01 | 0.31 |
|  |  |  |  | FarmerF5 | 0.01 | 0 | 0.02 | 0.06 |
|  |  |  |  | FarmerF6 | 0.01 | 0 | 0.02 | 0.12 |
| Shannon Diversity | Linear mixed model | Farm (Intercept) | 0.00 | Main_cropPangasius (Intercept) | 7.07 | 7.06 | 7.07 | 0 |
|  |  | Residual | 0.01 | Main_cropTilapia | 0.00 | 0.00 | 0.01 | 0.20 |
| Shannon Diversity | Linear mixed model | Farm (Intercept) | 0.00 | UpazilaJamalpur Sadar (Intercept) | 7.07 | 7.06 | 7.08 | 0 |
|  |  | Residual | 0.01 | UpazilaMuktagacha | 0 | -0.01 | 0 | 0.34 |
|  |  |  |  | UpazilaTarakanda | 0 | -0.01 | 0.01 | 0.92 |
| Shannon Diversity | Linear mixed model | Farmer (Intercept) | 0 | VillageBamunpara(Intercept) | 7.07 | 7.06 | 7.08 | 1 |
|  |  | Upazila (Intercept) | 0 | VillageFulbaria | 0 | -0.01 | 0.02 | 0.68 |
|  |  | Residual | 0.01 | VillageGopalpur | 0 | -0.01 | 0.02 | 1 |
|  |  |  |  | VillageKandulia | 0 | -0.02 | 0.01 | NaN |
|  |  |  |  | VillageMalotipur | -0.01 | -0.02 | 0.01 | 1 |
| Shannon Diversity | Linear mixed model | Farm (Intercept) | 0.00 | SeasonWinter (Intercept) | 7.07 | 7.06 | 7.07 | 0 |
|  |  | Residual | 0.01 | Seasonpost monsoon | 0 | -0.01 | 0.01 | 0.77 |

**Supplementary Table 2.**
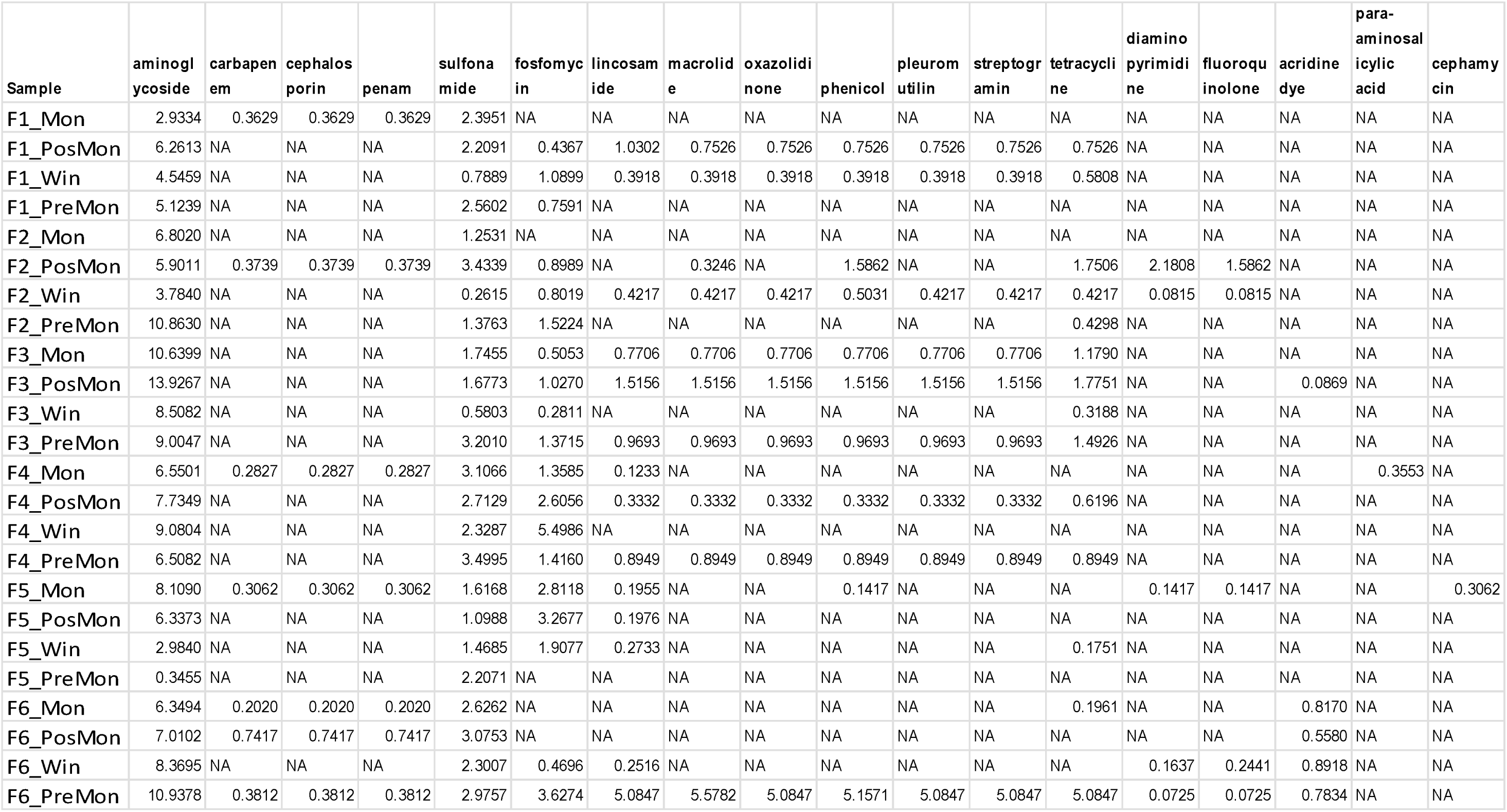
Sum of ARG abundance summarised by sample. ARGs are categorised by the antibiotic drug class they confer resistance to. ARG undance was calculated as transcripts per kilobase million (TPM).

| Sample | aminoglycoside | carbapenem | cephalosporin | penam | sulfonamide | fosfomycin | lincosamide | macrolide | oxazolidinone | phenicol | pleuromutilin | streptogramin | tetracycline | diaminopyrimidine | fluoroquinolone | acridine dye | para-aminosalicylic acid | cephamycin |
| --- | --- | --- | --- | --- | --- | --- | --- | --- | --- | --- | --- | --- | --- | --- | --- | --- | --- | --- |
| F1_Mon | 2.9334 | 0.3629 | 0.3629 | 0.3629 | 2.3951 | NA | NA | NA | NA | NA | NA | NA | NA | NA | NA | NA | NA | NA |
| F1_PosMon | 6.2613 | NA | NA | NA | 2.2091 | 0.4367 | 1.0302 | 0.7526 | 0.7526 | 0.7526 | 0.7526 | 0.7526 | 0.7526 | NA | NA | NA | NA | NA |
| F1_Win | 4.5459 | NA | NA | NA | 0.7889 | 1.0899 | 0.3918 | 0.3918 | 0.3918 | 0.3918 | 0.3918 | 0.3918 | 0.5808 | NA | NA | NA | NA | NA |
| F1_PreMon | 5.1239 | NA | NA | NA | 2.5602 | 0.7591 | NA | NA | NA | NA | NA | NA | NA | NA | NA | NA | NA | NA |
| F2_Mon | 6.8020 | NA | NA | NA | 1.2531 | NA | NA | NA | NA | NA | NA | NA | NA | NA | NA | NA | NA | NA |
| F2_PosMon | 5.9011 | 0.3739 | 0.3739 | 0.3739 | 3.4339 | 0.8989 | NA | 0.3246 | NA | 1.5862 | NA | NA | 1.7506 | 2.1808 | 1.5862 | NA | NA | NA |
| F2_Win | 3.7840 | NA | NA | NA | 0.2615 | 0.8019 | 0.4217 | 0.4217 | 0.4217 | 0.5031 | 0.4217 | 0.4217 | 0.4217 | 0.0815 | 0.0815 | NA | NA | NA |
| F2_PreMon | 10.8630 | NA | NA | NA | 1.3763 | 1.5224 | NA | NA | NA | NA | NA | NA | 0.4298 | NA | NA | NA | NA | NA |
| F3_Mon | 10.6399 | NA | NA | NA | 1.7455 | 0.5053 | 0.7706 | 0.7706 | 0.7706 | 0.7706 | 0.7706 | 0.7706 | 1.1790 | NA | NA | NA | NA | NA |
| F3_PosMon | 13.9267 | NA | NA | NA | 1.6773 | 1.0270 | 1.5156 | 1.5156 | 1.5156 | 1.5156 | 1.5156 | 1.5156 | 1.7751 | NA | NA | 0.0869 | NA | NA |
| F3_Win | 8.5082 | NA | NA | NA | 0.5803 | 0.2811 | NA | NA | NA | NA | NA | NA | 0.3188 | NA | NA | NA | NA | NA |
| F3_PreMon | 9.0047 | NA | NA | NA | 3.2010 | 1.3715 | 0.9693 | 0.9693 | 0.9693 | 0.9693 | 0.9693 | 0.9693 | 1.4926 | NA | NA | NA | NA | NA |
| F4_Mon | 6.5501 | 0.2827 | 0.2827 | 0.2827 | 3.1066 | 1.3585 | 0.1233 | NA | NA | NA | NA | NA | NA | NA | NA | NA | 0.3553 | NA |
| F4_PosMon | 7.7349 | NA | NA | NA | 2.7129 | 2.6056 | 0.3332 | 0.3332 | 0.3332 | 0.3332 | 0.3332 | 0.3332 | 0.6196 | NA | NA | NA | NA | NA |
| F4_Win | 9.0804 | NA | NA | NA | 2.3287 | 5.4986 | NA | NA | NA | NA | NA | NA | NA | NA | NA | NA | NA | NA |
| F4_PreMon | 6.5082 | NA | NA | NA | 3.4995 | 1.4160 | 0.8949 | 0.8949 | 0.8949 | 0.8949 | 0.8949 | 0.8949 | 0.8949 | NA | NA | NA | NA | NA |
| F5_Mon | 8.1090 | 0.3062 | 0.3062 | 0.3062 | 1.6168 | 2.8118 | 0.1955 | NA | NA | 0.1417 | NA | NA | NA | 0.1417 | 0.1417 | NA | NA | 0.3062 |
| F5_PosMon | 6.3373 | NA | NA | NA | 1.0988 | 3.2677 | 0.1976 | NA | NA | NA | NA | NA | NA | NA | NA | NA | NA | NA |
| F5_Win | 2.9840 | NA | NA | NA | 1.4685 | 1.9077 | 0.2733 | NA | NA | NA | NA | NA | 0.1751 | NA | NA | NA | NA | NA |
| F5_PreMon | 0.3455 | NA | NA | NA | 2.2071 | NA | NA | NA | NA | NA | NA | NA | NA | NA | NA | NA | NA | NA |
| F6_Mon | 6.3494 | 0.2020 | 0.2020 | 0.2020 | 2.6262 | NA | NA | NA | NA | NA | NA | NA | 0.1961 | NA | NA | 0.8170 | NA | NA |
| F6_PosMon | 7.0102 | 0.7417 | 0.7417 | 0.7417 | 3.0753 | NA | NA | NA | NA | NA | NA | NA | NA | NA | NA | 0.5580 | NA | NA |
| F6_Win | 8.3695 | NA | NA | NA | 2.3007 | 0.4696 | 0.2516 | NA | NA | NA | NA | NA | NA | 0.1637 | 0.2441 | 0.8918 | NA | NA |
| F6_PreMon | 10.9378 | 0.3812 | 0.3812 | 0.3812 | 2.9757 | 3.6274 | 5.0847 | 5.5782 | 5.0847 | 5.1571 | 5.0847 | 5.0847 | 5.0847 | 0.0725 | 0.0725 | 0.7834 | NA | NA |

**Supplementary Table 3.** Presence and abundance of the recA gene in gene calls from metagenomic assemblies found using a recA hidden Markov model at 1e^-50^ cut-off

| Sample | RecA genes | TPM |
| --- | --- | --- |
| F1_Mon | 577 | 352.19 |
| F1_PosMon | 540 | 298.29 |
| F1_Win | 521 | 261.55 |
| F1_PreMon | 321 | 217.33 |
| F2_Mon | 641 | 299.68 |
| F2_PosMon | 636 | 354.42 |
| F2_Win | 499 | 393.54 |
| F2_PreMon | 638 | 357.17 |
| F3_Mon | 783 | 457.16 |
| F3_PosMon | 774 | 486.69 |
| F3_Win | 527 | 383.24 |
| F3_PreMon | 579 | 329.53 |
| F4_Mon | 627 | 361.41 |
| F4_PosMon | 511 | 308.38 |
| F4_Win | 395 | 267.55 |
| F4_PreMon | 417 | 270.35 |
| F5_Mon | 587 | 315.63 |
| F5_PosMon | 437 | 288.86 |
| F5_Win | 515 | 248.50 |
| F5_PreMon | 331 | 228.78 |
| F6_Mon | 547 | 343.82 |
| F6_PosMon | 282 | 259.98 |
| F6_Win | 395 | 306.72 |
| F6_PreMon | 343 | 267.57 |
| **AVERAGE** | 517.63 | 319.10 |

**Supplementary Table 4.**
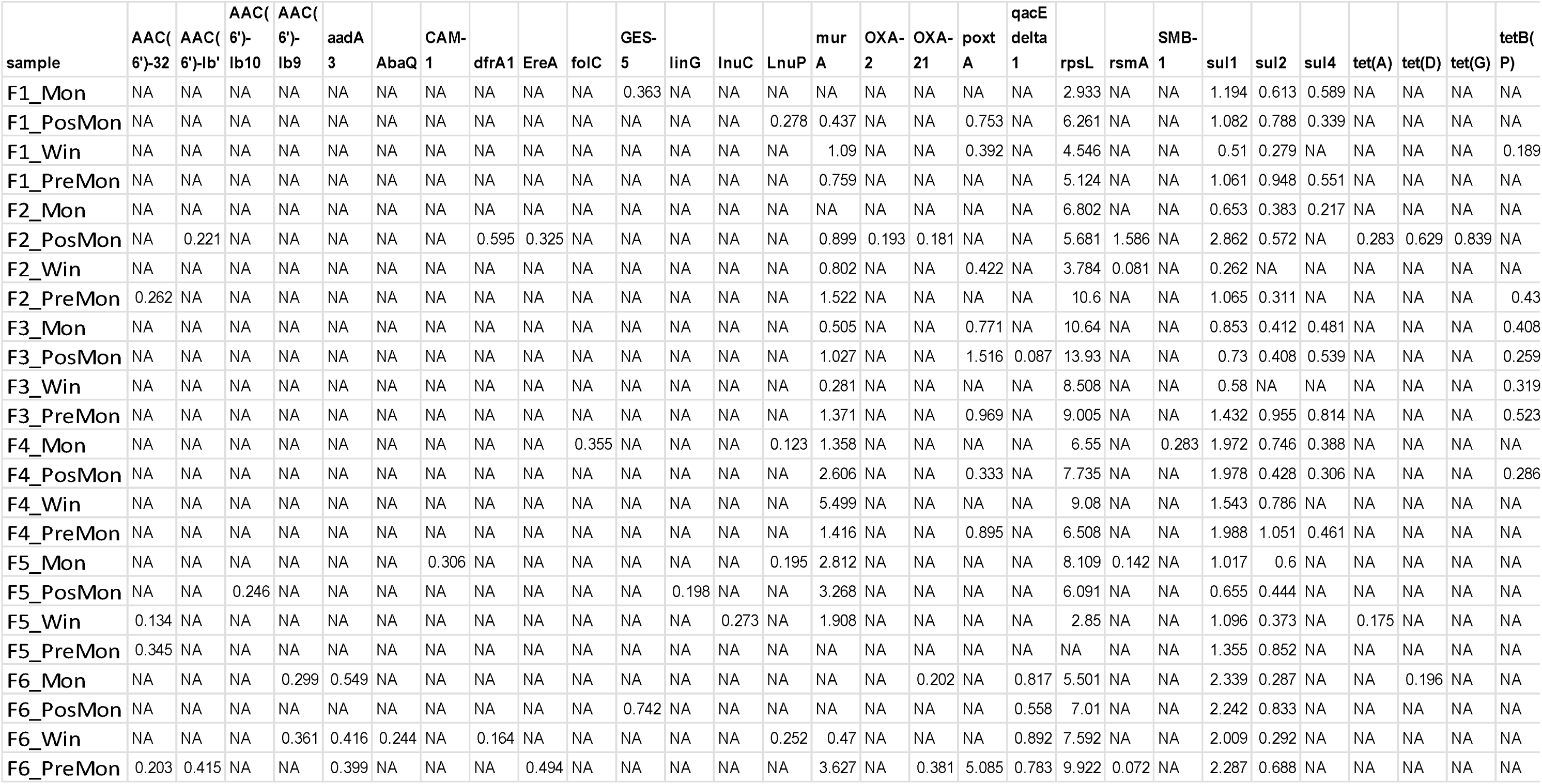
Sum of ARG abundance summarised by sample. ARG abundance was calculated as transcripts per kilobase million (TPM).

## CRediT authorship contribution statement

**Ashley Bell:** Software, Formal analysis, Investigation, Data Curation, Writing - original draft, Writing - review & editing, Visualization. **Kelly Thornber:** Conceptualization, Resources, Writing - Review & Editing, Supervision. **Dominique L. Chaput:** Methodology, Investigation, Data Curation, Writing - Review & Editing. **Neaz Al Hasan:** Investigation, Writing - Review & Editing. **Md. Mehedi Alam:** Investigation, Writing - Review & Editing. **Mohammed Mahfujul Haque:** Investigation, Writing - Review & Editing, Resources, Funding acquisition. **Jo Cable:** Conceptualization, Writing - Review & Editing, Supervision. **Ben Temperton:** Conceptualization, Methodology, Validation, Resources, Writing - Review & Editing, Supervision. **Charles R. Tyler:** Conceptualization, Resources, Writing - Review & Editing, Supervision, Project administration, Funding acquisition.

## Acknowledgements

The authors would like to acknowledge the use of the University of Exeter High-Performance Computing (HPC) facility in carrying out this work. We would also like to thank Jamie McMurtrie for his advice in data visualisation and analysis, and Sanjit Debnath for his knowledge in rural Bangladesh finfish aquaculture.

